# Csde1 binds transcripts involved in protein homeostasis and controls their expression in erythropoiesis

**DOI:** 10.1101/203497

**Authors:** Kat S Moore, Nurcan Yagci, Floris van Alphen, Nahuel A Paolini, Rastislav Horos, Ntsiki M Held, Riekelt H Houtkooper, Emile van den Akker, Alexander B Meijer, Peter A.C. ‘t Hoen, Marieke von Lindern

## Abstract

Expression of the RNA-binding protein Csde1 (Cold shock domain protein e1) is strongly upregulated during erythropoiesis compared to other hematopoietic lineages. In the severe congenital anemia Diamond Blackfan Anemia (DBA), however, Csde1 expression is impaired. Reduced expression of Csde1 in healthy erythroblasts impaired their proliferation and differentiation, which suggests an important role for Csde1 in erythropoiesis. To investigate the cellular pathways controlled by Csde1 in erythropoiesis, we identified the transcripts that physically associate with Csde1 in erythroid cells. These mainly encoded proteins involved in ribogenesis, mRNA translation and protein degradation, but also proteins associated with the mitochondrial respiratory chain and mitosis. Crispr/Cas9-mediated deletion of the first cold shock domain of Csde1 affected RNA expression and/or protein expression of Csde1-bound transcripts. For instance, protein expression of Pabpc1 was enhanced while *Pabpc1* mRNA expression was reduced indicating more efficient translation of Pabpc1 followed by negative feedback on mRNA stability. Overall, the effect of reduced Csde1 function on mRNA stability and translation of Csde1-bound transcripts was modest. Clones with complete loss of Csde1, however, could not be generated. We suggest that Csde1 is involved in feed-back control in protein homeostasis and that it dampens stochastic changes in mRNA expression.

## Introduction

RNA binding proteins (RBP) regulate transcript stability and translation. RBPs can cooperate with protein complexes of the general mRNA translation machinery, or with protein complexes that control mRNA location and/or degradation. Deregulated expression of such RBPs affects protein synthesis from a set of transcripts particularly dependent on that specific RBP. This has been referred to as an RNA regulon ^1^. The RNA regulon may be cell-type specific, because it depends on the available transcriptome in these cells.

Defects in ribosomal proteins that are involved in ribosome biosynthesis cause Diamond Blackfan Anemia (DBA), a severe congenital anemia ^2^. Interestingly, mutations in the erythroid transcription factor Gata1 also cause a DBA-like anemia ^3,4^, which suggests that at least a part of the Gata1 target genes form a RNA regulon that is very sensitive to reduced ribosome availability ^5^. Reduced expression of DBA-related ribosomal proteins in erythroblasts impairs translation of IRES (internal ribosomal entry site) containing transcripts ^6^. This could be due to the less competitive nature of IRES-mediated recruitment of ribosomes to a transcript compared to cap-dependent ribosomal recruitment. In addition, the aberrant expression of RNA binding proteins could enhance the effect of reduced ribosome availability on mRNA translation. An RBP that is repressed in erythroblasts of DBA patients is Csde1 (Cold shock domain protein e1)^6^.

Csde1 was first described as Unr (upstream of N-ras) in *Drosophila melanogaster* ^7^. It binds an AG-rich domain in the 3’UTR of *Msl* (*Male sex lethal*) and suppresses translation ^8,9^. Its expression is highly conserved across species and Csde1 binds its own IRES to repress translation in mammalian cells ^10^. Csde1 also enhances IRES-dependent translation of select transcripts, such as *Apaf1 (apoptotic peptidase activating factor 1)* ^11^ and *Cdk11B* (*cyclin dependent kinase 11B*) ^12,13^. Overall, the role of Csde1 in control of mRNA stability and translation may be diverse as it binds transcripts through distinct sites ^14,15^.

The variable effect of Csde1 on its bound transcripts may be explained by the RNA context and by the associated proteins that control the RNA binding affinity. Csde1 cooperates with PTB and hnRNP C1/C2 to control IRES-dependent translation of *Apaf1* and *Cdk11B*, respectively ^8,13,16^, it cooperates with DDX6 and miRNA in translational repression and P-body assembly *^17^*, and it acts together with Pabp to control mRNA translation and stability through elements in the 3’-UTR of specific transcripts ^18,19^. Thus, the molecular effects of Csde1 may be diverse. Increased expression of Csde1 has been associated with melanoma and breast cancer^20,21^. Analysis of Csde1-bound transcripts in melanoma implied Csde1 in control of metastasis as it increased the expression of, for instance, Vimentin ^20^.

Csde1 expression is much increased in erythroblasts compared to other hematopoietic cells, and reduced expression in primary mouse erythroblasts impaired their proliferation and differentiation similar to knock down of ribosomal proteins ^6^. Thus, Csde1 controls important erythroblast functions, that must differ from previously described functions such as sex specification or migration during metastasis. To identify its function in erythropoiesis, we aimed to identify the transcripts that are bound by Csde1 in erythroblasts, and to evaluate the effect Csde1 on transcript stability and translation. Csde1-bound transcripts mainly encoded proteins involved in protein homeostasis, ranging from ribosome biosynthesis, translation, to protein degradation. In addition, Csde1 bound transcripts encoding proteins involved in mitosis. Protein homeostasis and mitosis are affected in DBA. Csde1 also reduced translation of several ribogenesis factors, and increased translation from reduced Pabpc1 transcript levels. Overall, we suggest that the function of Csde1 is involved in feed-back control during protein homeostasis and that it may dampen stochastic changes in gene expression.

## Results

### Pull down of Csde1-bound transcripts

To identify mRNA transcripts bound by Csde1, we expressed *in vivo* biotinylated Csde1 in erythroid cells. MEL cells expressing the prokaryotic biotin ligase BirA were transfected with a *Csde1* construct tagged with the recognition site of BirA ^6^. RNA-protein complexes containing Biotag-Csde1 were precipitated on streptavidin beads (figure 1A). Prior to precipitation of Csde1, cell lysates contained similar levels of endogenous Csde1 and biotag-Csde1. As expected, streptavidin beads specifically associated with biotag-Csde1 (figure 1A). The biotagged version of Csde1 manifests as a double band at a marginally higher molecular weight ^6^. As Csde1 binds the IRES in its own transcript, we used conditions that enriched for binding to Csde1 mRNA, using *Bag1, Tbp (TATA-binding protein)* or *18S* rRNA as background signals (figure 1B).

**Figure 1.**
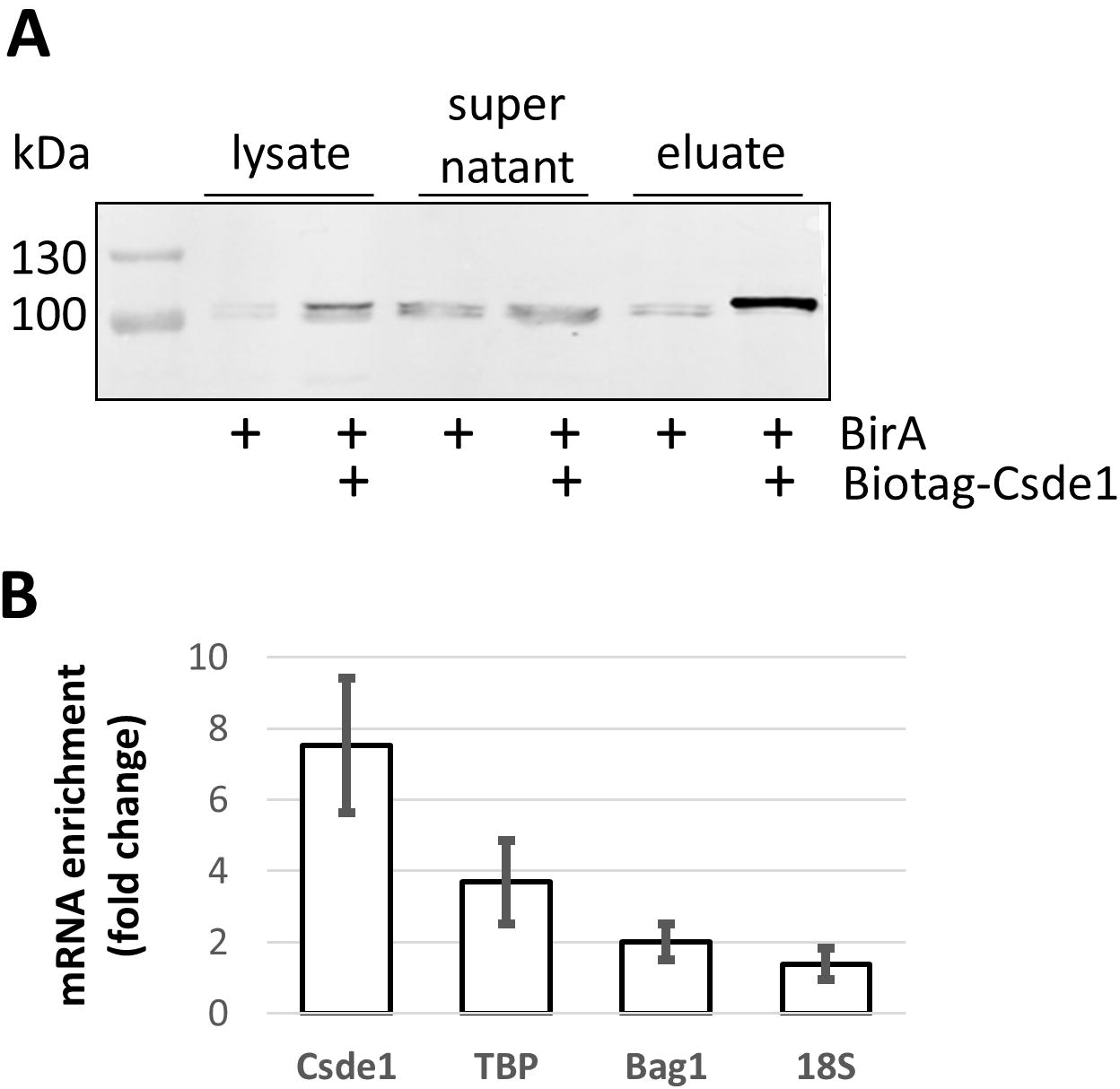
Purification of Csde1 containing RNPcomplexes. **(A)** MEL cells expressing BirA, or expressing BirA plus biotag-Csde1 were lysed, and incubated with streptavidin beads. Supernatant and beads were collected, beads were washed and eluted. Western blot with fractions was stained with anti-Csde1 antibody. Size markers are indicated in kD. As both endogenous and the biotagged Csde1 are expressed, the absence of a double band in the pulldown eluate indicates the strength of the biotin-streptavidin interaction. **(B)** RNA was isolated from eluates and tested for expression of *Csde1, Tbp, Bag1* and *18S* RNA by Q-PCR. The fold-change enrichment of the transcripts on streptavidin beads incubated with biotag-Csde1 lysate was calculated compared to pull downs from BirA MEL cells (error bars indicate SD, n=3).

### Identification of Csde1-bound transcripts

Streptavidin binding protein/RNA complexes were harvested from the cytoplasmic lysate of BirA expressing MEL cells that did or did not co-express biotag-Csde1. We isolated and processed RNA from three independent samples for sequence analysis with Illumina MiSeq; one sample was sequenced separately together with one control. Principle component analysis (PCA) discriminated the samples on sequence run in PC1, whereas PC2 discriminated the transcripts pulled down in biotag-Csde1/BirA expressing MEL cell lysates from the transcripts harvested from BirA control cell lysates (figure 2A). To identify the transcripts that are significantly enriched in biotag-Csde1-RNA complexes, we analyzed the results with DESeq2. Both significantly enriched and depleted transcripts were detected in Csde1-biotag/BirA MEL cells compared to BirA MEL cells, with a clear skewing towards enriched transcripts, as is to be expected during a pulldown (figure 2B, supplemental table S-II). The depleted transcripts represent the small set of constitutively biotinylated proteins (34 and supplemental table S-II) and supplemental table S-II). We selected transcripts enriched with a Benjamini-Hochberg false discovery rate (FDR) <0.05. This yielded a list with 292 unique transcripts enriched in Csde1-protein complexes (supplemental table S-III). Transcripts assigned to pseudogenes were included in this list because they may have a regulatory function by quenching RBP and miRNA.

**Figure 2.**
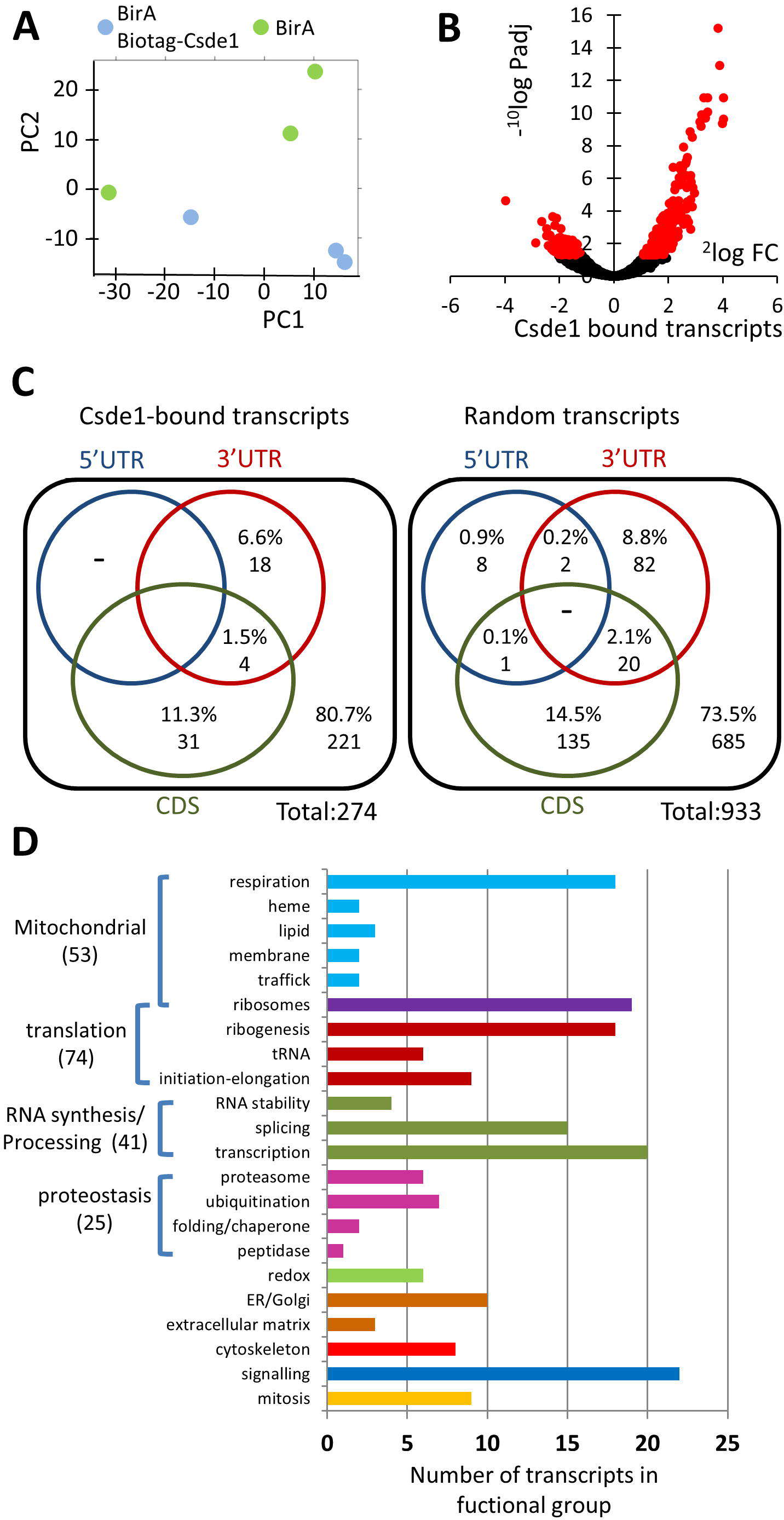
Identification of Csde1-bound transcripts. RNA was isolated from independent streptavidin bead eluates derived from MEL cells expressing BirA alone (N=3), or BirA plus biotag-Csde1 (N=3). RNA was sequenced, and differential abundance was analyzed by Deseq2. **(A)** Principle component analysis (green: BirA; blue BirA plus biotag-Csde1). **(B)** lOlog(P-adjusted) plotted against 2log(fold change) of differentially enriched transcripts. Red: FDR<0.05 **(C)** At FDR<0.05 we identified transcripts from 274 unique genes (pseudogenes excluded) that associated with Csde1. Potential Csde1 binding sites ([A/G]_5_AAGUA[A/G], or [A/G]_7_AAC[A/G]_2_) are indicated (percentage and total number) for the 5’UTR (blue), protein coding region (CDS; red) and 3’UTR (green); for the 274 Csde1-bound transcripts, and for almost 1000 control transcripts. **(D)** Each transcript was assigned a unique label that best represented its function. The number of transcripts within a specific function are depicted (overarching functions in the same color). Transcripts previously reported by Wurth et al. as Csde1 target genes, identified by iCLIP in melanoma cells are indicated in black^20^.

Previous studies using SELEX identified Csde1 binding sites as [A/G]_5_AAGUA[A/G], or [A/G]_7_AAC[A/G]_2_ ^35^. We performed a search for either of these consensus sites with a custom Python script using Biopython ^33^. They were present in 20% of Csde1-bound transcripts (60 out of 274, we excluded double counting on pseudogenes) versus 26% in random transcripts (248 out of 685) (figure 2C, supplemental table III). Not only the presence, but also the distribution of putative binding sites between the protein coding region (13% and 16%) and the 3’UTR (8% and 11%) in Csde1-bound and random transcripts was comparable (figure 2C). Thus, the presence of a consensus Csde1 binding site as determined by SELEX in transcripts is not predictive for Csde1 binding in erythroblasts. This may be explained by the conflicting finding that Csde1 binds a 6nt motif ([C/G/U]AAG[AUG]A), as identified by iCLIP ^20^. Due to its short length, this sequence is found ubiquitously among all detected transcripts (data not shown), making it unsuitable for *in silico* analysis.

To identify the cellular processes that may be regulated by Csde1, we classified the transcripts according to functional groups (e.g. transcription, translation, mitochondrial function; figure 2D, supplemental table IV). To determine whether cellular processes are significantly enriched, we used Overrepresentation Analysis (ORA) with GeneTrail2 ^32^; figure 2D, supplemental table IV). Mitochondrion and mitochondrial respiration were highly enriched among the cellular component and biological process GO-terms, respectively (53 hits). This includes mitochondrial ribosomes and ribosome association (n=23), the respiratory chain (n=19), lipid synthesis (n=3), heme synthesis (n=3), mitochondrial membrane (n=3) and transport of proteins to mitochondria (n=2) (figure 2D). Abundantly represented were processes involved in mRNA translation (n=74), including those associated with mitochondrial ribosomes (n=23), ribosome biogenesis (n=28), tRNA modifying enzymes (n=9) and mRNA translation initiation and elongation (n=14) (figure 2D). Also enriched were transcripts which affect protein synthesis via mRNA splicing (n=16) and mRNA stability (n=4). Csde1 targets additionally affect protein ability via maintenance of protein folding (n=4), and via the activity of peptidases (n=2), ubiquitinases (n=9) and the proteasome (n=10). The centrosome and control of mitosis were also significantly enriched terms (n=11). Recent studies identified Csde1 targets in melanoma cells and *Drosophila melanogaster*^20,36^. Comparison showed that 53 of the 274 transcripts we identified as Csde1-associated transcripts were also identified as a Csde1 target in melanoma cells using iCLIP ^20^. These common transcripts encoded proteins that act in all cellular processes, but were particularly abundant in control of translation and ribogenesis (figure 2D, supplemental table S-IV).

### Generating a model with reduced Csde1 expression

To investigate the role of Csde1 in expression and translation of Csde1-associated transcripts, we aimed to reduce the expression of Csde1 via shRNA transduction. Csde1 protein expression was clearly reduced in MEL cells targeted for shRNA-mediated Csde1 knockdown compared to SH002 control vector transduction (data not shown). However, analysis via mass spectrometry revealed that lentiviral transduction *per se* induced Csde1 expression. Therefore, shRNA-mediated knockdown merely reduced Csde1 to levels comparable to those measured in parental MEL cells, rather than representing a true reduction in Csde1 expression versus the untransduced state.

As an alternative to lentiviral knock-down, we used Crispr-Cas9 ^27^ to generate deletions in Csde1 in MEL cells. Knock down of Csde1 in primary cells abrogated their proliferation and differentiation ^6^. Therefore, we aimed for an in-frame deletion to remove the first cold shock domain, which causes a 20-fold reduction in RNA binding affinity ^35,37^. Guide RNAs in exon 3 (NM_144901.4), just downstream of the AUG start codon, and in exon 4 were transfected into MEL cells (figure 3A). Single cells were sorted by FACS (fluorescence assisted cell sorting) from the brightest top 5% of GFP-expressing cells. Selected clones were subsequently tested by PCR and Western blot for the intended deletion. This yielded two clones with an out-of-frame Csde1 deletion (*Del*; shown are clones D1, D2), three hypomorph clones with the intended in-frame deletion (*Hm*; shown are clones H1, H2), and multiple clones with a mono-allelic out-of-frame deletion in addition to a wt allele (heterozygote deletions, *Het*, indicated as clones C1, C2) (figure 3B). It is noteworthy that the control-transfected clones expressed Csde1 protein similar to parental MEL cells. Clones H1 and H2 expressed a shorter Csde1 protein isoform, as expected (figure 3C). Surprisingly, clones D1 and D2 were expected to lose Csde1 expression, but anti-Csde1 antibody recognized proteins of 70-75 kDa (figure 3C).

**Figure 3.**
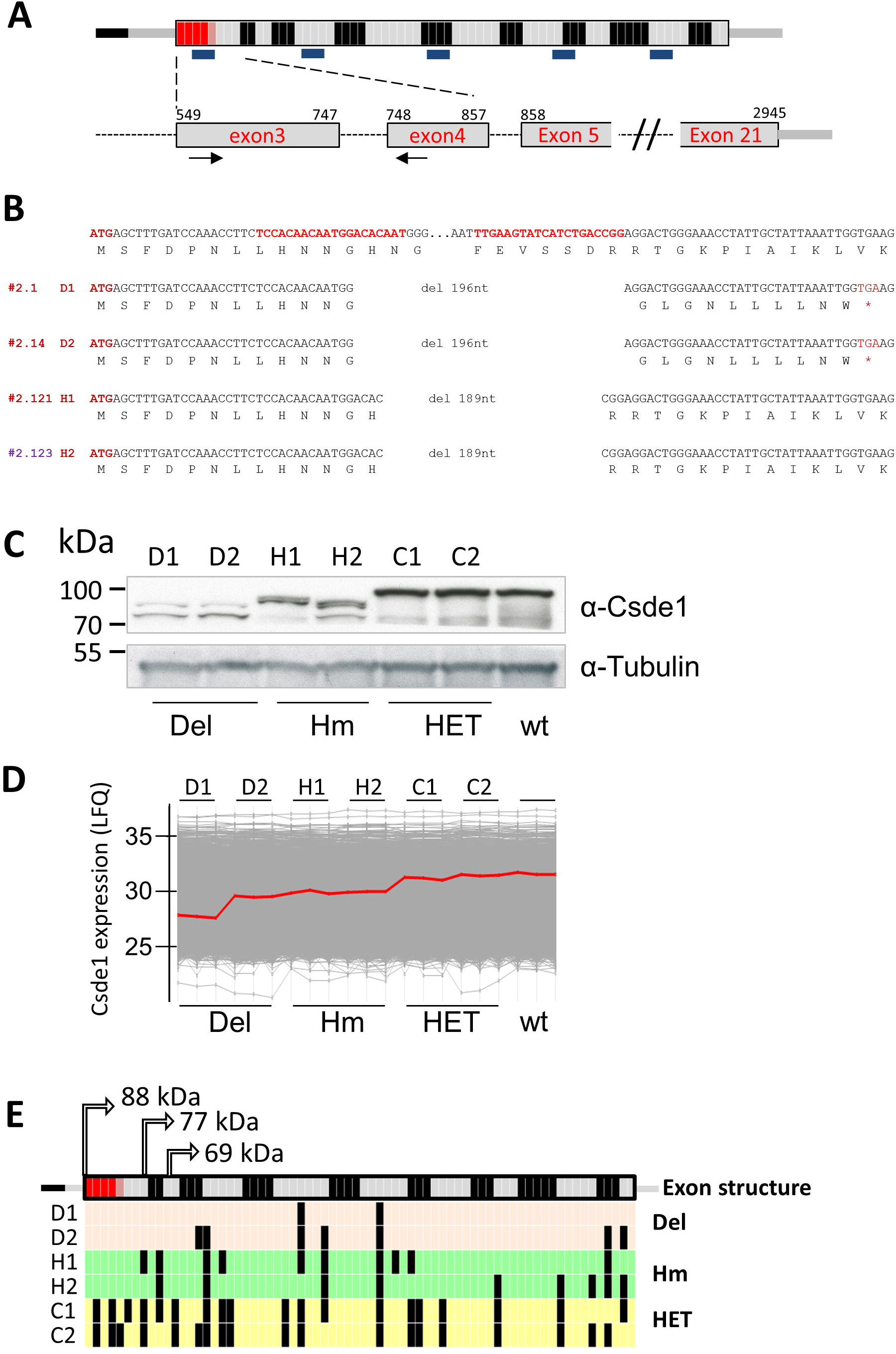
Deletion of the 1^st^ cold shock domain of Csde1 using Crispr/Cas9. **(A)** Cartoon of the Csde1 transcript. Grey and black represent subsequent exons. Small squares represent individual tryptic peptides (exons and peptides in arbitrary size). Tryptic peptides were assigned to the exon that contributes most. The methionine start codon locates at the start of exon 3. The position of five cold shock domains is indicated by short bars below the transcript. Numbers on the zoom in on exons 3-5 indicate the nucleotide position of NM_144901.4. Guide RNAs are indicated with arrows, the red tryptic fragments are affected by the deletion. **(B)** The top sequence shows the guide RNAs in red, in their sequence context, and the amino acids below the codons. Below the sequence that reveals the precise deletion for 4 clones. **(C)** Western blot stained for Csde1 and tubulin as loading control. D1 (2.1) and D2 (2.14) represent out-of-frame deletions (Del), H1 (2.121) and H2 (2.123) in-frame deletions of the 1^st^ cold shock domain (Hm). HET are control clones that harbor 1 deleted (out-of-frame) allele and 1 wt allele. **(D)** Mass spectrometry analysis with MaxQuant and Perseus yields a cartoon in which the protein expression of all proteins is indicated as LFQ by a grey line. Csde1 expression is indicated with a red line. **(E)** For Csde1, we identified 33 unique tryptic peptides. Black lines indicate tryptic fragments that were detected by mass spectrometry. Alternative AUG start codons that may explain truncated proteins are indicated by arrows, the size of the encoded protein in kDa.

Protein lysates of clones D1, D2, H1, H2, C1, C2 and wt MEL cells were submitted to mass spectrometry with label-free quantification. MaxQuant was used for peptide identification and quantification, and expression of Csde1 was verified ^24^. In accordance with Western blot data, Csde1 expression was similar in control clones and parental wt MEL cells, and reduced in clones H1 and H2 (figure 3D). Intriguingly, the out-of-frame deletion in the N-terminus of Csde1 should have abrogated Csde1 expression in clones D1 and D2, but Csde1 peptides were detected by mass spectrometry (figure 3D). Thus, both the Western blot and mass spectrometry suggest the expression of a shorter form of Csde1 in clones D1 and D2. To assess which part of the protein is expressed, the detected Csde1 peptides in the various clones were mapped, which showed that 33 of 72 predicted tryptic peptides were detected in parental MEL lines and in heterozygous deletion clones. (figure 3E). Several Csde1 peptides were detected in the clones that were assumed to lack Csde1 (figure 3E). These peptides were located downstream of the deletion, and downstream of a potential in frame start codon. The number of peptides is too small to draw conclusions on the start site. In addition, the sequence of the predicted novel N-terminal tryptic peptides is too short to be specific. Ribosome footprints deposited in the GWIPS database (http://gwips.ucc.ie/),^38^ indicate translation of several small uORFs in the 5’UTR of Csde1, which may facilitate leaky scanning and expression of smaller Csde1 isoforms^39^; supplemental figure 1). The bi-allelic out-of-frame deletion in *Csde1* did not affect the proliferation of MEL cell clones D1 and D2 (data not shown). This is in contrast to the observed abrogation of proliferation in primary erythroblasts after Csde1 knockdown ^6^. This strongly suggests that selective pressure induced leaky scanning and translation initiation downstream of the Crispr/Cas9 deletion on clones D1 and D2. Outgrowth of these clones was likely due to alternative translation initiation and expression of an N-terminally truncated Csde1 protein.

Because the transcripts associated with Csde1 were largely enriched for mRNAs encoding mitochondrial ribosomal proteins, and proteins of the mitochondrial respiratory chain, we investigated whether Csde1 expression and function may control mitochondrial activity and capacity. Csde1 was expressed in mitochondria, although at low expression levels (supplemental figure S2A). Mitochondrial respiration of Hm and Del clones was measured by Seahorse technology, but not altered in the Del and Hm clones compared to control clones (supplemental figure S2B).

### Protein and RNA expression in Del and Hm Csde1 mutant clones

Binding of Csde1 to target transcripts may affect transcript stability and/or translation. Therefore, we examined both protein and RNA expression in the N-terminally truncated/deleted clones D1, D2, H1, H2, and in control clones C1, C2 and wt MEL parental cells.

The peptide profiles of the two Del clones, the Hm clones, the control clones, and the parental BirA-MEL cells were subjected to cluster analysis. A total of 985 proteins were differentially expressed proteins in any of the clones (ANOVA, S0 cutoff 0.4, FDR cutoff 5%) (supplemental table V). Hierarchical clustering using the Perseus computational platform for proteomics analysis yielded a heatmap that showed a striking difference between the Hm clones and the Del clones ^25^. (figure 4A). The N-terminal truncation due to leaky scanning and translation initiation downstream of the deletion results in a protein with smaller size compared to the in-frame-deletion. It is possible that the Del clones were selected for efficient leaky scanning, which may have altered translation regulation on a larger scale.

**Figure 4.**
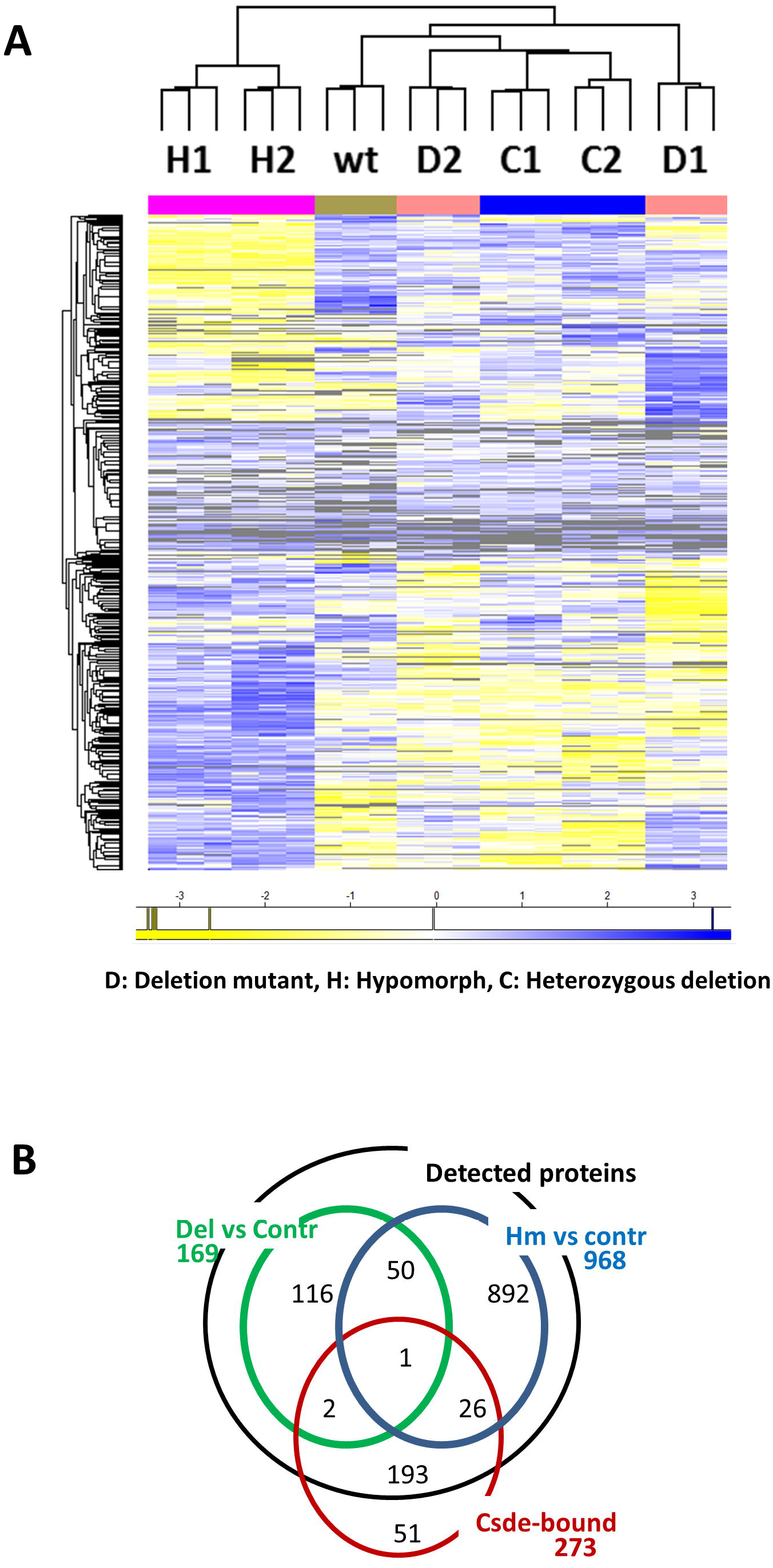
Proteome of Csde1 deletion mutants. Total MEL cell lysates of 2 Hm clones (H1, H2), 2 Del clones (D1, D2), 2 clones with a heterozygous out-of-frame deletion (C1, C2), and the parental MEL cells (wt) were analysed by mass spectrometry. Three lysates were analysed for each clone. Data were analysed by MaxQuant and Perseus **(A)** Proteins differentially regulated between Hm and control clones (C1, C2 and wt), or between Del and control clones were subjected to Pearson clustering, (blue: upregulated, yellow: downregulated, grey: not detected). Proteins and their Z-score in the order of this heatmap are listed in supplemental table S-V. **(B)** The overlap between genes encoding proteins differentially expressed between Del and control clones (169, green circle); between Hm and control genes (968, blue circle) and genes encoding Csde1-bound transcripts of which proteins were detected by mass spectrometry *(222*, red circle) (FDR < 0.05).

For 222 of the 274 Csde1-associated transcripts, we detected peptides at least in all Del or Hm clones, or in all control clones (excluding non-translated pseudogenes). There was little overlap, however, between differentially expressed proteins and Csde1-associated transcripts (figure 4B, supplemental table S-V) suggesting that the differentially regulated proteins are primarily secondary targets of Csde1-controlled pathways. The only Csde1-associated transcript corresponding to a differentially expressed protein in both Del and Hm clones versus control clones is *Pabpc1* (*PolyA binding protein C1*; supplemental table V, see discussion).

RBPs such as Csde1 control RNA stability and translation. To judge the role of Csde1, we generated RNA expression profiles, sequencing poly-adenylated RNA of the Del, Hm, and control MEL cells. Following normalization, we assessed the expression of *Csde1* in these clones. Expression of *Csde1* mRNA was reduced in clones D1, D2, H1 and H2, but also in clone C1, compared to C2 and parental MEL cells (figure 5A). The Crispr/Cas9-induced in frame deletion is not expected to affect transcript levels. The out-of-frame deletion, however, is expected to cause nonsense-mediated decay (NMD) due to splice factors residing at the many downstream splice junction sites ^40^. The Hm clones may carry an inframe deleted allele (figure 3B), and an out-of-frame deletion that is lost due to NMD. This could explain the lower expression in the Hm clones compared to the parental MEL cells. Importantly, the Del clones were expressed at similar levels as Hm clones, the expression was not lost due to NMD. This is in accordance with the proposed leaky scanning and translation of a shorter Csde1 protein isoform, which would protect the *Csde1* transcript from NMD. Of note, the first in frame AUG codon downstream of the deletion occurs in exon 4, between the deletion and the first splice junction that could give rise to NMD (figure 3E).

**Figure 5.**
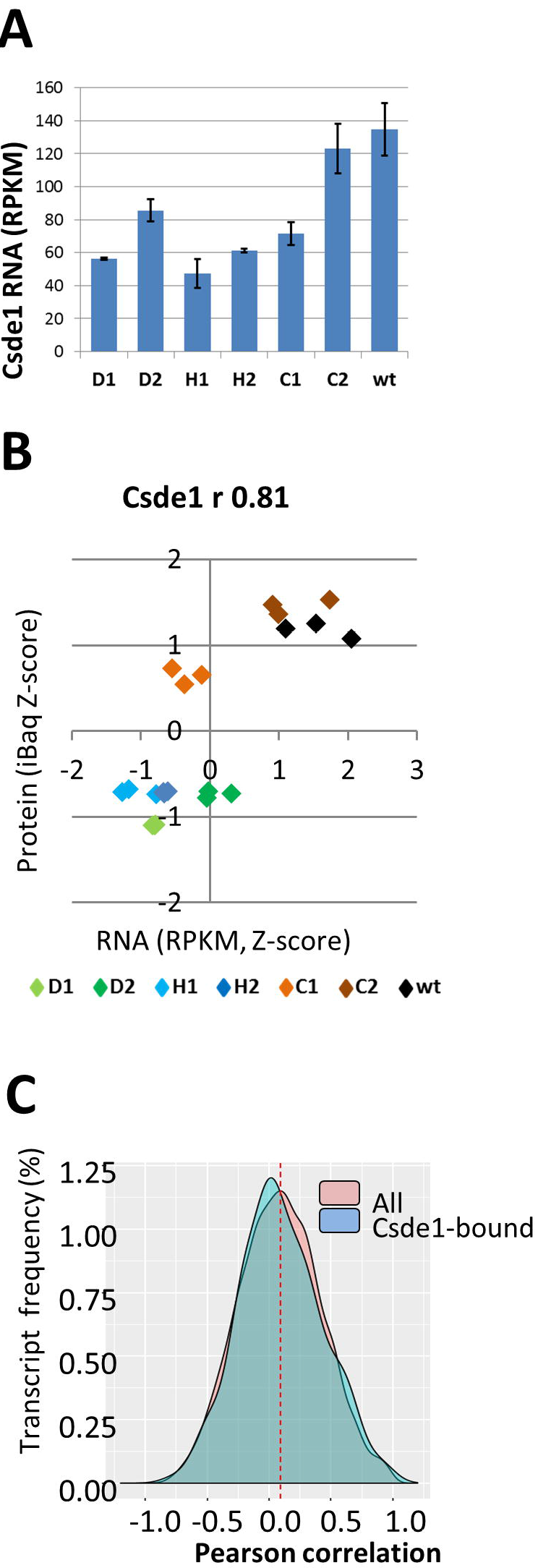
The correlation between protein and mRNA expression. RNA was isolated in triplicate from 2 Hm clones (H1, H2), 2 Del clones (D1, D2), 2 clones with a heterozygous out-of-frame deletion (C1, C2), and the parental MEL cells (wt), and polyAdenylated RNA was sequenced. RNA reads were normalized and calculated as RPKM (reads per kilobase per million). **(A)** RNA expression of Csde1 in RPKM **(B)** The same cell isolates were used for RNA and protein analysis (figure 4). For each protein detected by mass spectrometry, the Pearson correlation (r) between protein (Z-scores of iBaq) and RNA (Z-scores of RPKM) was calculated for the 21 samples. The distribution of the correlation between −1 and +1 was plotted for all samples, and for transcripts bound by Csde1. **(C)** the correlation between protein and RNA was plotted for Csde (Z-scores, standard deviations from the mean). (Del clones: green; Hm clones: blue, HET control clones: orange-brown, wtMEL: black)

A likelihood ratio test (LRT) followed by analysis of deviance by clone (ANODEV) was used to identify transcripts which differed significantly in expression across all samples. These transcripts were combined in a heatmap for all differentially expressed genes (supplemental figure S3; supplemental Table S-VI). Strikingly, the RNAseq heatmap is very different from the proteome heatmap (figure 4A). In a PCA, the samples are poorly separated (supplemental figure S4a).

### Correlation between protein and RNA expression of Csde1-bound transcripts

Correlation between mRNA abundance and protein expression is widely understood to be poor ^41^, though the exact strength of correlation depends widely on the study and methods employed. Findings generally indicate correlations between 0.2 and 0.4 across a variety of organisms and tissue types ^41–47^, though some authors have argued that such studies may underestimate the strength of correlation between mRNA and proteins ^48^. The absence of strong correlation between mRNA and protein expression levels can be due to a number of post-transcriptional and post-translational factors, including the presence of regulatory proteins acting as translational moderators, such as Csde1. We therefore investigated whether loss of Csde1 alters the strength of correlation between mRNA and protein abundancy. To this end, we calculated RPKM (Reads Per Kilobase of transcript per Million mapped reads) to quantify RNA expression, and normalized iBAQ values to quantify protein expression (supplemental table S-VII). For Csde1 itself, this clearly visualized that both RNA and protein expression of Csde1 were reduced in the MEL clones D1, D2, H1, H2 compared to low RNA and intermediate protein expression in clone C1 and increased RNA and protein expression in clones H2 and wt MEL cells (figure 5B). Looking broadly at all transcripts, the Spearman rank correlation coefficient between ^10^log(RPKM) and ^10^log(iBAQ) varied between 0.52 and 0.56 for all samples (supplemental table S-VIII). This is similar to the observed Pearson correlation of 0.59 between RNA and protein expression in two mouse hematopoietic cell lines ^46^. Surprisingly, the correlation coefficient differed only marginally for Csde1-bound transcripts compared to the transcript pool as a whole.

We also calculated the Pearson correlation coefficient between the Z scores of RNA (RPKM) and protein (iBAQ) for individual transcripts across conditions (supplemental table S-IX). The distribution of correlation coefficients showed that the strength of correlation can vary widely. The modal correlation coefficient between protein and RNA expression was 0.1, but there was no apparent difference between Csde1-bound transcripts (222) and random transcripts (6400; figure 5C).

Although Csde1 does not influence the correlation between protein and mRNA for all its associated transcripts, it is possible that Csde1 may regulate the balance of protein expression for a sub-selection of target transcripts. We focused on the Csde1-bound transcripts and plotted RNA expression against protein expression. Of particular interest is that Csde1 target transcripts associated with protein degradation displayed distinct patterns of RNA and protein expression. Protein expression levels of the proteasome subunits *Psme1* (r 0.96) and *Psmc3* (r −0,78) were highly correlated and anti-correlated, respectively. Both RNA and protein expression of clones D1, H1 and H2 were increased for *Psme1*, whereas for Psmc3, protein levels in these clones decreased while RNA levels increased (figure 6A,B). *Npepps* (*Aminopeptidase Puromycin Sensitive*) transcript levels of Del and Hm clones showed the same variation as control clones, but protein levels were increased, whereas RNA/protein relation was scattered for *Fkbp11* (*Fk506 Binding Protein 11*) (figure 6C,D).

**Figure 6.**
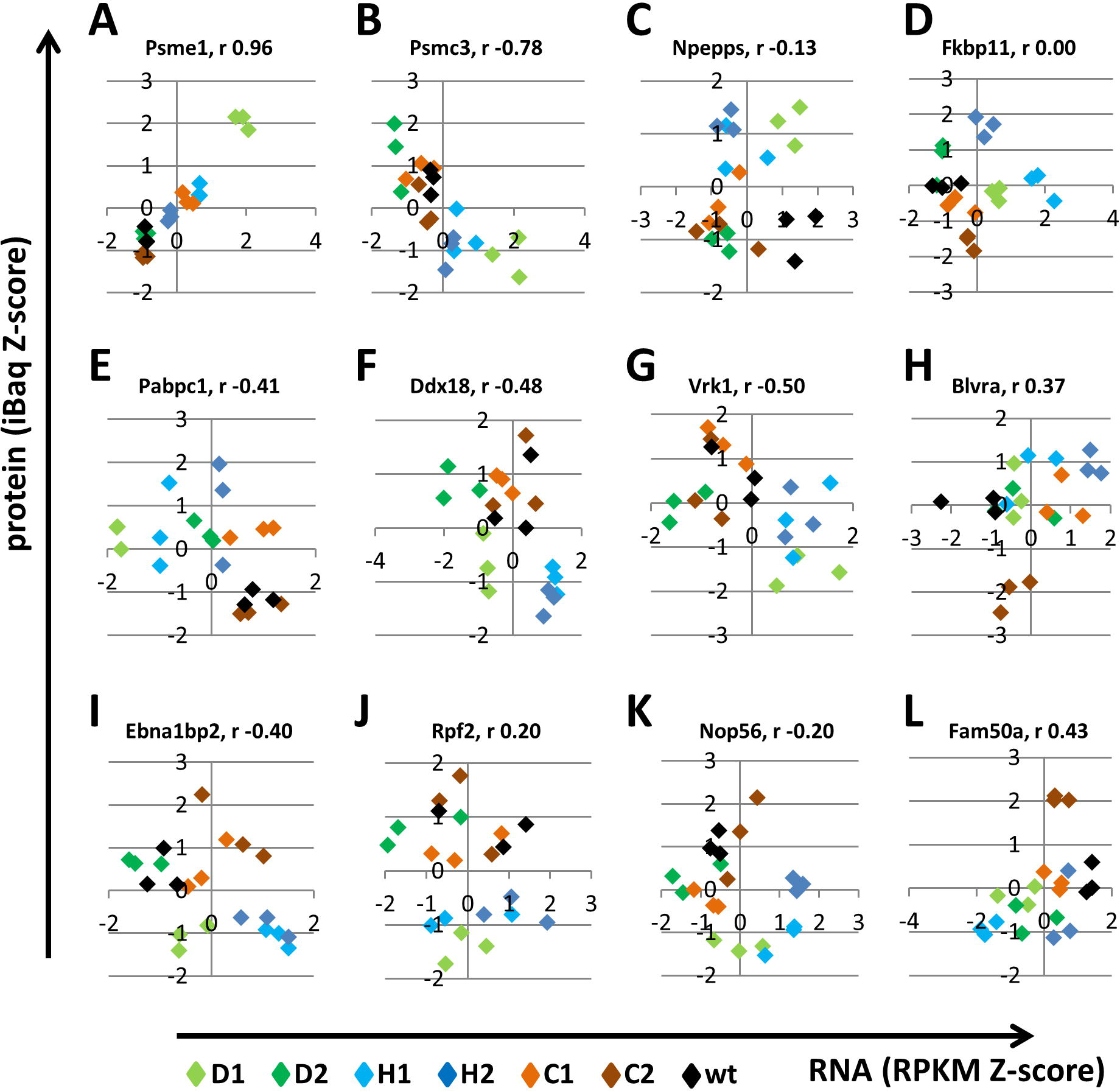
The correlation between protein and mRNA expression for few Csde1-bound transcripts. The correlation between protein and RNA was plotted for the indicated genes (Z-scores of iBaq and RPKM; see legend to figure 5). (Del clones: green; Hm clones: blue, HET control clones: brown, wtMEL: black)

Pabpc1 was the only protein encoded by a Csde1-bound transcript that was significantly changed in the Del and Hm clones compared to control clones (figure 4B). Interestingly, Pabpc1 protein expression was increased whereas *Pabpc1* RNA expression was reduced in Del and Hm clones compared to control clones, which hints to feedback control (figure 6E, see discussion). In contrast, the splicing factors *Ddx18 (Dead Box Polypeptide 18)* and *Vrk1 (Vaccinia Related Kinase 1)* displayed lower protein expression from increased transcript levels in Del and Hm clones (figure 6F,G). *Csde1* was identified as being poorly translated in DBA, a ribosomopathy. Therefore, it is striking that three nucleolar proteins involved in ribogenesis (*Ebna1bp2, Ebna1 Binding Protein 2; Rpf2, Ribosome Production Factor 2; Nop56, nucleolar protein 56*) showed reduced protein expression in clones D1, H1 and H2 relative to controls whereas mRNA levels were similar (figure 6I,J,K). Additional transcripts with a clear segregation of Del and Hm clones versus control clones were *Blvra (Biliverdin Reductase A)*, *Fam50a (Family With Sequence Similarity 50, Member A)*, *Aldh2 (Aldehyde Dehydrogenase 2)*, *eIF3h (eukaryotic initiation factor 3h)*, *Eprs (Glutamyl-Prolyl-Trna Synthetase)* and *Rps8 (Ribosomal protein S8)* (figure 6H,L, supplemental figure S4B). In conclusion, reduced Csde1 expression/function caused either increased or reduced protein expression. This suggests that the function of Csde1 may be determined by the protein complex in which it acts on the fate of the bound transcript.

### Discussion

Csde1 is a RNA binding protein that is strongly upregulated during erythropoiesis. Its expression is reduced in erythroblasts of patients suffering from Diamond Blackfan Anemia, a severe congenital anemia due to impaired ribosome biogenesis. Csde1 is a critical protein in erythropoiesis, but the pathways controlled by Csde1 in erythropoiesis are unknown. We show that Csde1-bound transcripts in erythroblasts mainly encode proteins involved in ribogenesis, in mRNA translation and protein stability, and in mitochondrial function. Deletion of the N-terminal cold shock domain by Crispr/Cas9 resulted in truncated proteins due to in-frame deletions, or due to translation initiation downstream of out-of-frame deletions. The expression levels of mRNA and/or protein of multiple Csde1-bound transcripts were consistently changed in Csde1-mutated cells compared to control clones. Pabpc1 protein levels are increased, whereas the encoding mRNA is decreased. In contrast, the nucleolar ribogenesis factors Ebna1bp2, Rpf2, and Nop56 showed reduced protein expression in Del and Hm clones from comparable mRNA levels. The results suggest a general role for Csde1 in the regulation of transcript stability and translation, and a central role in protein homeostasis aimed at dampening stochastic changes in gene expression.

We identified Csde1 as a protein that is downregulated as a result of reduced ribosomes in DBA ^6^. This study reveals that Csde1 itself also controls ribogenesis and mRNA translation. Among the proteins involved in ribogenesis are Rpf2, Nop56 and Ebna1bp2 ^49,50^. *Nop56* and *Ebna1bp2* also interacted with Csde1 in melanoma cells ^20^. Expression of these proteins was reduced in Del and Hm clones. Rpf2 cooperates with Rpl5 and Rpl11 to incorporate 5S rRNA in the 60S ribosomal subunit. Of note, haploinsufficiency of RPL5 and RPL11 is a frequent cause of DBA. Nop56 is part of the Box C/D snoRNP complex and involved in rRNA methylation. Ebna1bp2 functions as a nucleolar scaffold protein. Two other proteins involved in ribosomal subunit maturation, Bola3 and Nol12, were bound by Csde1. However, their expression was too low for reliable detection by mass spectroscopy under the conditions used. Reduced efficacy of Csde1, and subsequent reduced expression of Rpf2, Nop56 and Ebna1bp2, is predicted to impair ribosome biogenesis ^51,52^. Further studies are needed to confirm the governing role of Csde1 on ribogenesis. The mechanism through which the reduction in Csde1 efficacy cooperates with haploinsufficiency of ribosomal proteins as observed in DBA is uncertain. It could further reduce ribogenesis, but it may also restore the balance between rRNA and ribosomal proteins and result in less p53 activation.

In the context of DBA, it is noteworthy that the identified Csde1-bound transcripts includes 11 transcripts involved in cell cycle regulation, all of which encode proteins that function in mitosis. This includes centrosome-regulating proteins Aurkaip1 (Aurora kinase interacting protein), Ccdc77 (Coiled-Coil Domain Containing 77), Spc24 (Ndc80 Kinetochore Complex Component), and Tubgcp2 (Tubulin, Gamma Complex Associated Protein 2), whereas Actn4 (Actinin Alpha 4), Ccdc124 (Coiled-Coil Domain Containing 124), Cdkn3 (Cyclin-Dependent Kinase Inhibitor 3) and Tubgcp2 (Tubulin, Gamma Complex Associated Protein 2) control cytokinesis. Csde1 is known to control translation of Cdk11B (PITSLRE) during mitosis in HEK293 cells, which is IRES-driven and requires cooperation with hnRNP C1/C2 ^13^. Expression of Cdk11 was not detected in erythroblasts using mass spectroscopy. We speculated previously that the occurrence of binucleated cells in erythroblasts that lack Rps19 (DBA-derived or upon knock down) is due to dysregulation of the centrosome or cytokinesis ^6^. As polyribosome recruitment of Csde1 is diminished upon loss of Rps19, it is possible that disruption of Csde1 impairs cytokinesis via deregulation of these target proteins, resulting in a binucleated phenotype. This provides a possible mechanism for Csde1’s role in DBA.

### mRNA stability and translation of Csde1-bound transcripts

The Del and Hm clones displayed reduced *Pabpc1* mRNA expression, and increased Pabpc1 protein expression. Actually, *Pabpc1* was the only Csde1-bound transcript that was significantly deregulated in Del and Hm clones. *Pabpc1* is an important target because it enhances both mRNA stability and translation in general. Pabpc1 binds the polyA tail of transcripts, which protects them from deadenylation and subsequent degradation ^53^. Pabpc1 simultaneously binds eIF4G scaffold of the cap-binding complexes which loops the polyA tail of a transcript to the start of the transcript and is supposed to enhance reassociation of ribosomal subunits for a new round of translation ^54^. In addition, Pabpc1 is involved in ribonucleoprotein complexes that regulate the stability or translation of distinct transcripts. Csde1/UNR was shown to cooperate with Pabp in the combined regulation of mRNA stability and translation of several transcripts using distinct mechanisms ^18,19,55–57^. Most importantly, Csde1/UNR forms a complex with Pabpc1 and Imp1 that binds an adenine-rich autoregulatory sequence (ARS) in the 5’UTR of *Pabp*. The ability of mutated ARS sequences to bind the trimeric UNR/Imp/Pabp complex correlated with their repression of *Pabp* translation ^58^, though it was not shown in their report that reduced Csde1/UNR expression affected *Pabp* expression. The existence of this trimeric complex predicts that loss of Csde1 function will increase protein expression of Pabpc1, as we observed. Increased Pabp levels will mitigate the repression, which will limit the effects of reduced Csde1 function. It implies that regulation of *Pabpc1* by Csde1 may constitute a feedback loop that dampens fluctuations in mRNA translation.

The interaction of Csde1/Unr with Pabpc1 was also shown to bind to sequences in other transcripts to control mRNA stability and translation. For instance, Csde1-Pabp binds a sequence within the coding region of *cFos* which enhances translation-dependent degradation ^19^. This mechanism predicts increased mRNA levels upon reduced Csde1 expression, which is observed for *Vrk1* and other negatively correlated Csde1 targets. Overall, however, the changes in the proteome correlate better to the clone genotype (Del vs Hm vs control) than the changes in mRNA translation (compare figures 4 and S4). Csde1 seems to control protein expression more than mRNA expression.

Whereas the Csde1-bound transcripts and encoded proteins show relatively small differences in expression levels, the overall proteome is vastly different. Strikingly, the Hm clones are much alike, but very different from the Del clones. This phenomenon is most likely due to clonal variation in translation initiation and efficiency. It was surprising that mass spectrometry detected Csde1 peptides in MEL clones harboring a Cas9-induced out-of-frame deletion. The deletion we aimed at started within the first tryptic peptide of Csde1 encoded by exon 3, and spans 5 tryptic peptides to end within the first tryptic peptide encoded by exon 4. Three of these peptides are detected in the control clones, but not in the clones harboring a deletion. Moreover, we only detect smaller proteins in de MEL clones harboring a deletion. These proteins most likely arose from alternative start codon recognition. The 5’UTR of Csde1 harbors several translated uORFs, which enables skipping of the first AUG start codon ^38,39^,and translation initiation at downstream start codons in a favorable Kozak consensus sequence. These are present at the end of exon 4 (the same exon that is targeted for the deletion), and in exon 6 (figure 3E). Complete loss of Csde1 is not compatible with embryonic development^59^, and Csde1 has have a pLI score of 1.0 in the Exome Aggregation Consortium database ^60^, indicating an extreme intolerance for Loss of Function (LoF) mutations. One recent study has found that cold shock domains 2 and 4 are the only cold shock domains required (out of 5) for translational stimulation^56^, though there is also evidence to suggest that that all five cold shock domains contribute to the ability of Csde1 to stimulate translation, especially from IRESs ^37^. The data strongly suggest that MEL cells carrying an out-of-frame deletion underwent selection to maximize leaky scanning. This change in translation initiation efficiency will change the entire proteome, as approximately 50% of all transcripts carries an uORF, which renders protein expression dependent on the rate of leaky scanning (manuscript in preparation)^39,61,62^.

The direct targets of Csde1 in erythroblasts are mainly ribogenesis factors, translation factors and protein degradation factors. The changes in expression of these regulators can be expected to amplify the expression of some secondary targets. Alternatively, the example of Pabpc1 indicates that the role of Csde1 is to stabilize protein expression by dampening stochastic changes in protein expression. Ribosome biosynthesis, translation factors and protein degradation factors were highly enriched among direct Csde1 targets, which suggests that Csde1 is mainly important in protein homeostasis in erythroblasts. The recently published study on Csde1 targets in melanoma focused on the effect of Csde1 on migration and metastasis, but the network shown for melanoma cells, and the comparison of targets between the two studies indicated that Csde1 controls protein synthesis and stability also in melanoma ^20^. In the context of DBA, this may be a very important function. It is a riddle why the genotype-phenotype correlation in DBA is so poor. With the advance of molecular diagnostics, it became clear that some parents of DBA patients were carriers of the DBA mutation without showing any signs of the disease. Siblings with the same mutation may show widely different pathology. This suggests that haploinsufficiency of ribosomal proteins and reduced expression of, among others, Csde1 cause a destabilized protein expression profile that is easily disturbed, but can also maintain a healthy state. Loss of Csde1 as a stabilizing factor in protein expression may therefore contribute to the variability in disease penetrance.

## Materials and Methods

### Cell culture

Murine erythroleukemia (Mel) and HEK293T cells were cultured in RPMI, and DMEM respectively (Thermofisher), supplemented with 10% (vol/vol) fetal calf serum (FCS; Bodinco), glutamine and Pen-Strep (Thermofisher). Mel cells expressing BirA, or BirA plus biotag-Csde1 were described previously^6^. Cell number and size were determined using CASY cell counting technology (Roche).

### Lentivirus production and transductions

HEK293Ts were transfected with pLKO.1-puro lentiviral construct containing shRNA sequences for Csde1: TRCN0000181609 and a scrambled control shRNA: SHC002 (MISSION TRC-Mm 1.0 shRNA library; Sigma-Aldrich; available on the BloodWeb site), pMD2.G, and pSPAX.2 packaging plasmids (gift of T. van Dijk, Erasmus MC, Rotterdam, The Netherlands) using 0.5M CaCl2 and HEPES (Thermofisher). 72 hours after transduction, viral supernatant was harvested and concentrated using 5% w/v PEG8000 (Sigma). Mel cells were transduced with a multiplicity of infection of 3-5 and addition of 8 ug/mL of Polybrene (Sigma-Aldrich). Transduced cells were selected with 1 ug/ml puromycin 24 hours after transduction.

### Protein-RNA pulldown for Csde1

100 million Mel-BirA and Mel-BirA-Csde1-tag cells were subjected to RNA immunoprecipitation using the protocol described by Keene et al 2006^22^, with the following modifications. M-270 Dynabeads (Thermofisher) were utilized in a volume of 100μl per 100 million cells. The Dynabeads were blocked for 1 hour at 4°C in 5% chicken egg albumin and then washed 3× in ice-cold NT2 buffer consisting of 50mM Tris-HCl (Sigma-Aldrich), 150mM NaCl (Sigma-Aldrich), 1mM MgCl_2_ (Thermofisher) and 0.05% NP40 (Sigma-Aldrich) prior to use. The beads were then resuspended in 850ul cold NT2, supplemented by 200U RNAse Out (EMD Bioscience), 400μM vanadyl ribonucleoside complexes (VRC, New England Biolabs) and 20mM EDTA (EM Science). Incubation was done for 2 hours at 4°C. The beads were then immobilized in a magnet rack and washed 5× with NT2 in 0.3M NaCl. At this point, the beads were split into a protein and an RNA fraction. The protein fraction was eluted via boiling in l× Laemmli buffer (Sigma-Aldrich) for 5 minutes. RNA fractions were purified using Trizol (Invitrogen), precipitated in isopropanol and washed in 75% ethanol.

### SDS-PAGE and Western blotting

Proteins were detected via SDS-PAGE and Western blotting as described in Horos et al, 2012 ^6^. Antibodies used were directed against Csde1 (NBP1-71915, Novus Biological), Actin (A3853, Sigma-Aldrich) and alpha Tubulin (ab4074, Abcam). Fluorescently labeled secondary antibodies for visualization with Odyssey were IRDye 680RD Donkey anti-Rabbit IgG (926-68073, Licor) and IRDye 800CW Donkey anti-Mouse IgG (925-32212, Licor).

### cDNA synthesis and qRT-PCR

cDNA was generated from 1μg RNA, using 1μg random primers (48190011, Invitrogen), 50U M-MLV reverse transcriptase (Invitrogen), 1mM dNTPs (Invitrogen) in M-MLV reverse transcriptase buffer (Invitrogen) supplemented with 5mM DTT (Thermofisher). The cDNA mix was heated for 45’ at 42°C and then for 3’ at 99°C.

Q-RT-PCR was performed as described in Horos et al, 2012^6^, with the following modifications. A master mix was created using 10μM of each primer, 12.5μl SYBR Green master mix (4309155, Applied Biosystems) and 5μl cDNA filled to a final concentration of 20μl. Primers can be found in Supplementary Table S-I.

### Mass spectrometry

Eluted peptides were processed as described by^23^. Samples were subjected to mass spectrometry using label-free quantification. All data was analyzed and processed with MaxQuant for peptide identification and quantification ^24^. Downstream statistical analysis was performed with Perseus v1.5.1.6 ^25^. All proteins matching the reverse database, potential contaminants, and those only identified by site were filtered out._To be considered for analysis, a protein had to be detectable within all triplicates of at least one clone. Prior to statistical testing, a log2 transformation was performed. Because failures to detect a given peptide is sometimes due to insufficient depth, missing values were imputed from the normal distribution with a width of 0.3 and a downshift of 1.8. These values were later de-imputed prior to visualization and production of the final tables. For multi-way ANOVA between CRISPR clones, an artificial within-group variance (S0) threshold of 0.4 was used ^26^. For two-way comparisons between groups, a *t*-test with a threshold of S0=0.5 was used. For all analyses, a Benjamini-Hochberg false discovery rate of < 0.05 was applied.

### Production of Csde1 CRISPR clones

Guide RNAs for Csde1 were designed using an online web tool from the Massachusetts Institute of Technology (http://crispr.mit.edu/). The probes were designed to target the sequences upstream and downstream of the first cold shock domain and selected on the basis of faithfulness to on-target activity (Supplementary Table S1). CRISPR clones were generated as described in Cong et al, 2013^27^. Briefly, the guide RNAs were ligated in the PX458 Cas9 expression vector and electroporated into Mel cells with the Amaxa SFcell line 4D-nucleofector XkitL. GFP positive cells were sorted using flow cytometry and deleted regions were checked using genotyping primers (Supplementary Table S1), Sanger sequencing and Western blotting.

### Seahorse

Mitochondrial respiration levels for Csde1 CRISPR clones were determined on Seahorse XF96 using the Seahorse XF Mito Stress Test kit. 24 hours prior to the assay, cells were seeded at a concentration of 150,000 in 500ul RPMI on a XF cell culture microplate. One hour before measurement, medium was replaced by DMEM (Sigma, D5030) containing 25 mM glucose (Sigma), 1 mM sodium pyruvate (Lonza), and 2 mM L-glutamine (Life technologies) and cells were incubated in a non-CO2 37C incubator. Basal oxygen consumption rate (OCR) was detected as an indicator of mitochondrial respiration. OCR was measured in response to injection of oligomycin (15 μM), FCCP (1μM), antimycin A (2.5 μM) and rotenone (1.25 μM). Experiments were performed in triplicate with 8 or 9 wells per experiment.

### RNA-sequencing

RNA-seq on RNA immunoprecipitated with Csde1 was performed by the Leiden Genome Technology Center (LGTC, Leiden), using library preparation following the template-switch protocol (Clontech) followed by Nexterea tagmentation. These samples were pooled together on one miSeq (Illumina) lane (2×l50bp, paired end). Sequence quality was checked using Fastqc (Babraham Bioinformatics). Quality control and trimming was performed using Trimmomatic with the following parameters: LEADING 20, TRAILING 20, SLIDINGWINDOW 4:20, MINLEN:50. We then used Tophat v2.0.9 ^28^ to align to mouse genome mm10 (Dec 2011) using the following parameters: library-type fr-unstranded, --mate-inner-dist 50, --mate-std-dev 20. The resulting bam files were sorted and indexed using samtools. Read count tables were produced using HTseq count^29^ in conjunction with a mouse mm10 gtf downloaded from the UCSC browser on 14 March 2014.

RNA expression by total mRNA sequencing from Csde1 CRISPR clones was performed by Novogene Co., LTD. Briefly, library preparation was performed using the NEB Next^®^ Ultra™ RNA Library Prep Kit and enriched using oligo(dT) beads. Isolated mRNA was fragmented randomly in fragmentation buffer, followed by cDNA synthesis using random hexamers and reverse transcriptase. After first-strand synthesis, a custom second-strand synthesis buffer (Illumina) was added with dNTPs, RNase H and Escherichia coli polymerase I to generate the second strand by nick-translation. The final cDNA library is ready after a round of purification, terminal repair, A-tailing, ligation of sequencing adapters, size selection and PCR enrichment. The complete library was sequenced using illumina HiSeq 2500 (2×150bp, paired end). Sequence quality was checked using Fastqc (Babraham Bioinformatics). Spliced Transcripts Alignment to a Reference (STAR,^30^) was used to align the sequences to the mouse mm10 genomic reference sequence, using the following parameters: --outFilterMultimapNmax 20, -- outFilterMismatchNmax 1, --outSAMmultNmax 1, -outSAMtype BAM SortedByCoordinates, quantMode GeneCounts, -outWigType wiggle, -outWigStrand Stranded, --outWigNorm RPM. A gtf file accessed from the UCSC genome browser on 11-Sept-2015 was passed to STAR using–sjdbGTFfile.

In both experiments, the read count tables were subjected to differential expression analysis with DESeq2 ^31^. DESeq2 implements a negative binomial generalized linear model to identify differential expressed/enriched transcripts. This method normalizes raw counts by adjusting for a size factor to account for discrepancies in sequencing depth between samples. The normalized counts are subsequently subjected to a Wald test with a Benjamini-Hochberg (FDR) correction for multiple testing, or a likelihood ratio test followed by analysis of deviance, in the case of multi-sample comparisons. DESeq2 also provides a function for principle component analysis (PCA). Additional visualizations were made using R packages ggplots, dheatmap and pheatmap. Overrepresentation Analysis (ORA) for GO-terms and pathways was performed on significant transcripts with GeneTrail2 ^32^.

### Identification of Csde1 binding sites in target transcripts

A custom Python script using Biopython was created to search transcripts for Csde1 binding sites^33^. Briefly, the script parses Genbank sequences into a Python dictionary and then scans the transcript for the presence of one of the known binding sites represented as regular expressions. Using the Genbank annotation, the script reports the location and exact sequence of the potential binding site(s). The script is available as supplemental information.

### Correlation of RNA and protein expression levels

To determine the degree to which RNA expression determines protein abundance, we calculated a Pearson correlation coefficient between the Z-scores from RNA sequencing and mass spectrometry data for each gene. Z-scores were calculated after normalization: In mass spectrometry, iBAQ values as determined via MaxQuant were normalized via a scaling factor calculated by dividing the sum of intensities from each sample by the intensity sum of a reference sample. RNA expression levels were normalized as reads per kilobase of transcript per million mapped reads (RPKM). When performing a sample-wise correlation between mRNA and protein expression, we utilized a Spearman rank correlation coefficient between ^10^log(RPKM) and ^10^log(iBAQ).

## Accession numbers

Original sequencing results have been deposited in the BioProject Database under project ID PRJNA378565. The mass spectrometry proteomics data have been deposited to the ProteomeXchange Consortium via the PRIDE partner repository with the dataset identifier PXD006358.

## Supplementary data

All supplementary tables are present as a single Excel file, Moore_supp_tables.xlsx, supplementary figures as Moore_supp_figs.ppt

## Acknowledgements

We want to thank Ben Nota for ICT support and Monika Wolkers for critical reading of the manuscript. Additionally, we would like to thank Klaske Thiadens, Fiamma Salerno, Franca di Summa and Aicha Ait Soussan for their support and expertise in the laboratory setting.

---

**Supplemental figure S1.** Single representative tracks were selected from the data set posted by De Klerk et al on GWIPS viz (http://gwips.ucc.ie/). All tracks are accessible at the website. Shown are exon 1 and 2 of Csde1 that represent the 5’UTR. Top tracks (blue) represent footprints obtained after 10 minutes treatment with Haringtonin to identify translation initiation sites. Lower tracks (red) are footprints obtained after stabilization of ribosomes on mRNA using cycloheximide to identify actively translated open reading frames. Green squares indicate translated upstream open reading frames (uORFs). For more information see De Klerk et al., 2015

**Supplemental figure S2. (A)** MEL cell lysate was fractionated in a cytoplamic fraction (C) and the remainder (-C). The remainder was fractionated in a nuclear fraction (N), a mitochondrial fraction (M) and the remainder (-M, -N). Fractions and total cell lysate (T) were assayed on Western blot for Csde1 expression. The Western blot was stained for Lamin B, Tubulin and Cytochrome C as controls for nuclear proteins, cytoplasmic proteins, and mitochondrial proteins, respectively. **(B)** Oxygen consumption rate (OCR) of Csde1 CRISPR clones versus wild-type, as measured by the Seahorse mitostress kit. OCR is measured in four phases. The first phase represents basal respiration, after which ATP-synthesis inhibitor oligomycin is injected to inhibit mitochondrial respiration. An injection of carbonyl cyanide p-trifluoromethoxy-phenylhydrazone (FCCP), which allows free migration of protons across the mitochondrial membrane, permits a measurement of maximum respiration. In phase 4, an injection of rotenone & antimycin A ends the mitochondrial overdrive state and returns the OCR to minimum activity. Values are corrected for cell input (ug DNA). Error bars represent the standard error of the mean (SEM).

**Supplemental figure S3.** RNA was isolated from MEL cells. Hm clones (H1, H2), Del clones (D1, D2), control clones (C1, C2) and parental MEL (wt). RNA was sequenced (polyA mRNA), and analysed. Three samples were analysed for each clone. Significant transcripts from ANODEV analysis were subjected to hierarchical clustering. (blue: upregulated, yellow downregulated).

**Supplemental figure S4. (A)** Principle component analysis of RNA-seq on Csde1 CRISPR clones. (Del clones: green; Hm clones: blue, HET control clones: orange-brown, wtMEL: black). **(B)** Correlation between mRNA and protein expression of select transcripts (Z-scores, standard deviations from the mean). (Del clones: green; Hm clones: blue, HET control clones: orange-brown, wtMEL: black)

